# Antagonistic regulation of salt and sugar chemotaxis plasticity by a single chemosensory neuron in *Caenorhabditis elegans*

**DOI:** 10.1101/2023.01.25.525622

**Authors:** Masahiro Tomioka, Yusuke Umemura, Yutaro Ueoka, Risshun Chin, Keita Katae, Chihiro Uchiyama, Yasuaki Ike, Yuichi Iino

## Abstract

The nematode *Caenorhabditis elegans* memorizes various external chemicals, such as ions and odorants, during feeding. Here we find that *C. elegans* is attracted to the monosaccharides glucose and fructose after exposure to these monosaccharides in the presence of food; however, it avoids them without conditioning. The attraction to glucose requires a left-sided ASE gustatory neuron called ASEL. ASEL activity increases when glucose concentration decreases. Optogenetic ASEL stimulation promotes forward movements; however, after glucose conditioning, it promotes turning, suggesting that after glucose conditioning, the behavioral output of ASEL activation switches toward glucose. We previously reported that chemotaxis toward sodium ion (Na^+^), which is sensed by ASEL, increases after Na^+^ conditioning in the presence of food. Interestingly, glucose conditioning decreases Na^+^ chemotaxis, and conversely, Na^+^ conditioning decreases glucose chemotaxis, suggesting the reciprocal inhibition of learned chemotaxis to distinct chemicals. The activation of PKC-1, an nPKC ε/η ortholog, in ASEL promotes glucose chemotaxis and decreases Na^+^ chemotaxis after glucose conditioning. Furthermore, genetic screening identified ENSA-1, an ortholog of the protein phosphatase inhibitor ARPP-16/19, which functions in parallel with PKC-1 in glucose-induced chemotactic learning toward distinct chemicals. These findings suggest that kinase–phosphatase signaling regulates the balance between learned behaviors based on glucose conditioning in ASEL, which might contribute to migration toward chemical compositions where the animals were previously fed.

**Author summary:** *Caenorhabditis elegans* responds to compounds that taste salty, bitter, sour, etc. However, its response to sweet compounds is unclear. Here, we show that *C. elegans* responds to glucose through a chemosensory neuron called ASEL. *C. elegans* avoids high concentrations of glucose and learns to approach glucose after feeding in the presence of high glucose, dependent on the action of ASEL. The ASEL neuron has been reported to respond to salt and promotes salt attraction after feeding in the presence of high salt. We find that the feeding-associated attractive responses to glucose and salt are antagonistic. When encountered with a mixture of salt and glucose during feeding, *C. elegans* changes its chemotactic response toward those chemicals according to the balance of each chemical in the mixture. *C. elegans* may memorize the concentrations of the chemical mixture during feeding and migrates to the chemical composition previously fed, which may promote opportunities obtaining food. Furthermore, we find that kinase–phosphatase signaling, which modulates neurotransmission, in ASEL is required for chemotaxis based on information processing of salt and glucose.

## Introduction

Animals sense information from complex environments and exhibit behavioral responses according to current and past situations. They receive multiple sensory cues, which are integrated in the nervous system to generate a repertoire of behaviors [1]. Molecular signaling pathways, such as ALK receptor tyrosine kinase and TGF-β signaling pathways, have been identified as regulators of the nervous system to control decision-making behaviors after sensing conflicting stimuli [2, 3]. The sensation of population density affects learning of sensory cues, such as odor chemotaxis learning and learned pathogen avoidance, by modulating neuropeptide secretion from chemosensory neurons through pheromonal signaling [4, 5]. Signaling derived from the gut affects decision-making and associative learning through insulin-like hormonal signaling [6, 7]. Although several signal transduction pathways and neural circuits that regulate information integration have been reported, our understanding of mechanisms of multisensory integration is limited.

*Caenorhabditis elegans* shows chemotactic responses toward multiple sensory stimuli, such as ions, temperature, and odorants, which dramatically change according to past experiences. They memorize concentrations of sodium chloride (NaCl) during feeding and are attracted toward those NaCl concentrations after feeding conditioning, thereby increasing the possibility of obtaining food [8]. A right-sided ASE gustatory neuron, ASER, senses NaCl and promotes attraction toward NaCl concentrations exposed to during feeding. Levels of diacylglycerol (DAG) are altered in the ASER axon according to changes in NaCl concentration, which in turn modulate the activity of PKC-1, an nPKC ε/η ortholog, to migrate to the NaCl concentration at which they were fed through changes in neurotransmission from ASER [9-11]. A left-sided ASEL neuron also contributes to NaCl chemotaxis. The ASEL neuron senses Na^+^ and promotes chemotaxis toward Na^+^ only after feeding in the presence of high NaCl concentrations; however, the mechanism of Na^+^ chemotaxis plasticity is largely unknown [12].

In addition to sensing the salty taste of NaCl, *C. elegans* senses a variety of tastes, including bitter, such as that from plant alkaloids, sour (acidic pH) and amino acids [13-15]. However, it is unclear whether and how it responds to sweet substances, such as mono- and disaccharides, whose sensing is beneficial for survival and has been extensively studied in mammals and insects [16]. Here, we show that *C. elegans* responds to monosaccharides, glucose and fructose. Worms show avoidance of these monosaccharides and are attracted toward the monosaccharides after feeding conditioning in the presence of high concentrations of monosaccharides. The ASEL neuron senses glucose and is required for glucose attraction after glucose conditioning. Similar to the action of ASER in NaCl chemotaxis plasticity, ASEL is activated when glucose concentration decreases and the behavioral output from ASEL is dramatically changed by glucose conditioning to promote high-glucose migration.

In mammalian taste buds, Receptor (type II) taste bud cells respond to a single taste stimulus from sweet, bitter, and umami tastes, whereas Presynaptic (type III) cells respond to all tastes, including sour and salty [17]. In *C. elegans*, taste compounds are mainly received by the sensory cilia of chemosensory neurons in the amphid sensory organ [18]. ASE, ASH, and ASK sensory neurons have been reported to respond to NaCl, bitter compounds, and amino acids, respectively [13, 14, 19]. Here, we examined the interaction between information of sugar (glucose) and salt (NaCl), which are both sensed by the same sensory neuron ASEL, in chemotaxis plasticity. Interestingly, glucose conditioning reduced Na^+^ attraction. Conversely, NaCl conditioning reduced glucose attraction, suggesting that information of glucose and Na^+^ are separately processed during conditioning and modulate chemotactic responses toward those chemicals in opposite directions. Furthermore, we investigated the mechanism of chemotaxis regulation for glucose and Na^+^ in different directions after glucose conditioning. This mechanism might underlie a survival strategy by which animals migrate toward precise locations at which they were previously fed using multiple chemicals.

## Results

### *C. elegans* is attracted toward glucose and fructose after feeding conditioning in the presence of the monosaccharides

A study reported that *C. elegans* is attracted to a region containing 75 mM glucose on agar plate [20]. To test the response of *C. elegans* to glucose, we prepared an agar plate with a glucose gradient (Supplemental Fig. 1A) and examined chemotaxis. Because the worms learn to be attracted to NaCl after feeding conditioning in the presence of high concentrations of NaCl [8] (Fig. 1A first block), we examined glucose chemotaxis after feeding conditioning in the presence of glucose. Worms were attracted to glucose after feeding conditioning with 20– 100 mM glucose for 5 h (Fig. 1A second block, B first block). By contrast, they showed significant avoidance of glucose after feeding conditioning without glucose (Fig. 1A second block). The significant attraction to glucose was not observed after starvation conditioning in the presence of glucose for 5 h, suggesting that the learned attraction to glucose requires both exposure to glucose and feeding experience during conditioning (Fig. 1B second block). Similarly, the worms showed significant attraction to fructose after fructose conditioning in the presence of food and significant avoidance of fructose without fructose conditioning (Fig. 1A third block). By contrast, the worms showed no substantial response to the disaccharide sucrose after sucrose conditioning in the presence of food (Fig. 1A fourth block). These data suggest that *C. elegans* is attracted toward the monosaccharides glucose and fructose after feeding conditioning in the presence of the monosaccharides. Hereafter, glucose was mainly used as a chemical cue to study the worms’ response to monosaccharides.

**Figure 1.**
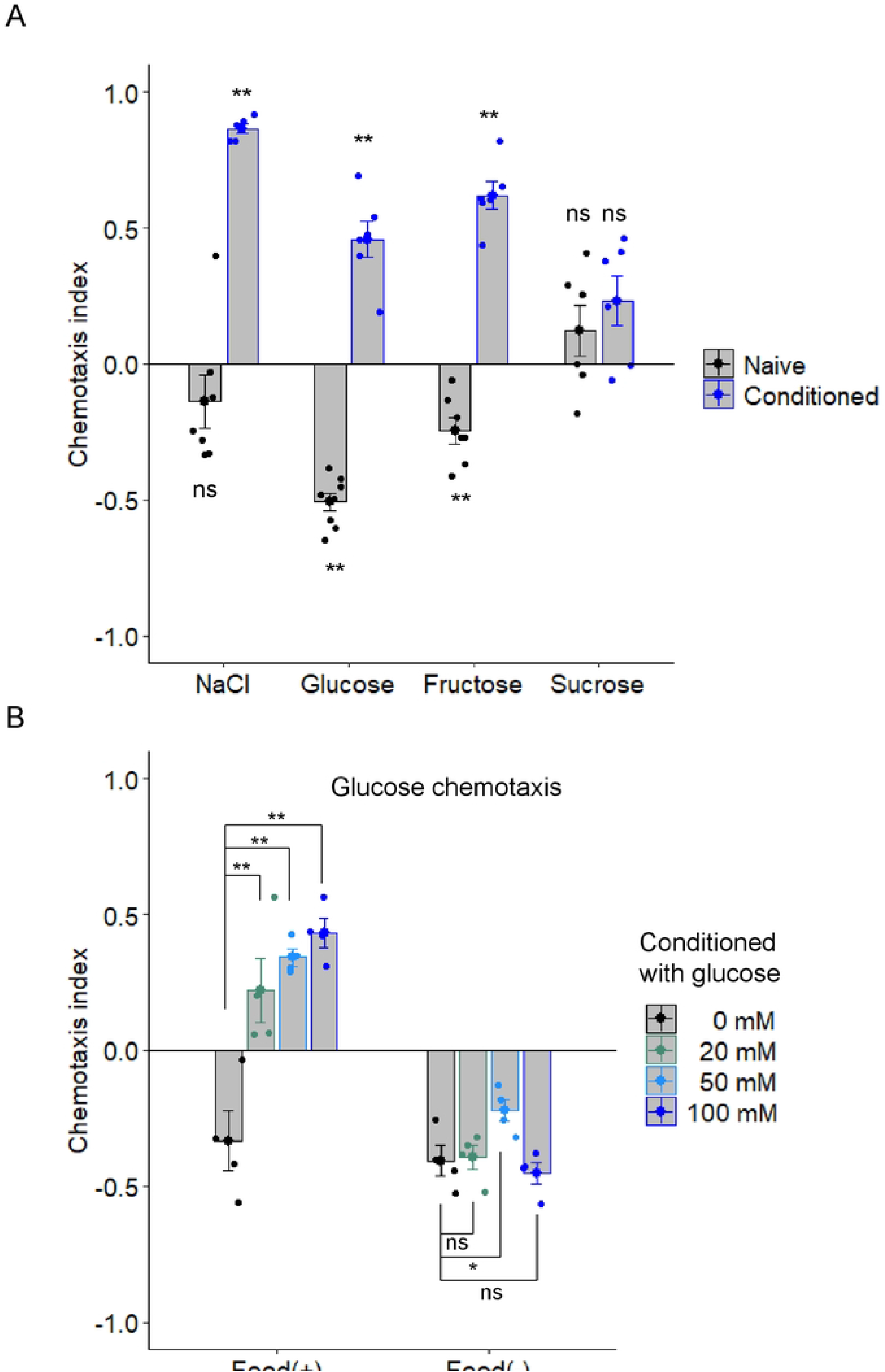
*C. elegans* memorizes the presence of monosaccharides during feeding. (A) After feeding conditioning with (“conditioned”) or without (“naïve”) a chemical indicated on the *x*-axis, chemotaxis to the chemical used for conditioning was tested. A chemotaxis index was determined according to the following equation: chemotaxis index = (N_A_ − N_B_) / (N_all_ − N_C_), where N_A_ and N_B_ are the number of worms in areas of high and low concentrations of chemical cues, respectively, N_all_ is the total number of worms on a test plate, and N_C_ is the number of worms in the area around the starting position. Positive and negative chemotaxis indices mean migration toward high and low concentrations of chemicals, respectively (see also Supplemental Figure 1A). Each blue or black dot represents a chemotaxis index calculated in each chemotaxis assay after conditioning with or without the chemicals indicated on the *x*-axis, respectively. n = 6–7 assays. (B) After conditioning with several concentrations of glucose in the presence (“Food (+)”) or absence (“Food (−)”) of food, chemotaxis to glucose was tested. n = 4 assays. Bars represent mean values; error bars represent SEM. One-sample, two-tailed *t*-test against zero value (A) or ANOVA followed by Dunnett’s *post hoc* test (B): \**P* < 0.05, \*\**P* < 0.01.

### A left-sided ASE gustatory neuron, ASEL, responds to glucose and is required for glucose attraction after glucose conditioning

ASE gustatory neurons play important roles in chemotaxis to water-soluble chemicals [21]. Left- and right-sided ASE neurons, ASEL and ASER, respectively, receive different ions and respond to increase and decrease, respectively, in concentrations of ions to promote attraction to those ions [19, 22]. We performed calcium imaging of the ASE neurons in response to changes in glucose concentration. ASEL, but not ASER, responded to a decrease in glucose concentration (Fig. 2A top, Supplemental Fig. 2). Calcium levels of ASEL remained high while glucose concentrations were low and rapidly ceased when glucose concentrations were increased. After glucose conditioning, ASEL was activated when glucose concentration decreased at the same level as that without conditioning (Fig. 2A top, B left). By contrast, at low glucose concentration, the calcium level was significantly higher after glucose conditioning than that without conditioning (Fig. 2A top, B right), implying differences in ASEL properties between worms with and without glucose conditioning. The ASEL neuron of an *unc-13* mutant, in which synaptic transmission is largely disrupted because of exocytosis defects, responded to the decrease in glucose concentration at the same level as that in the wild type (Fig. 2A bottom, B), suggesting that ASEL is activated following direct sensing of ambient glucose concentrations.

**Figure 2.**
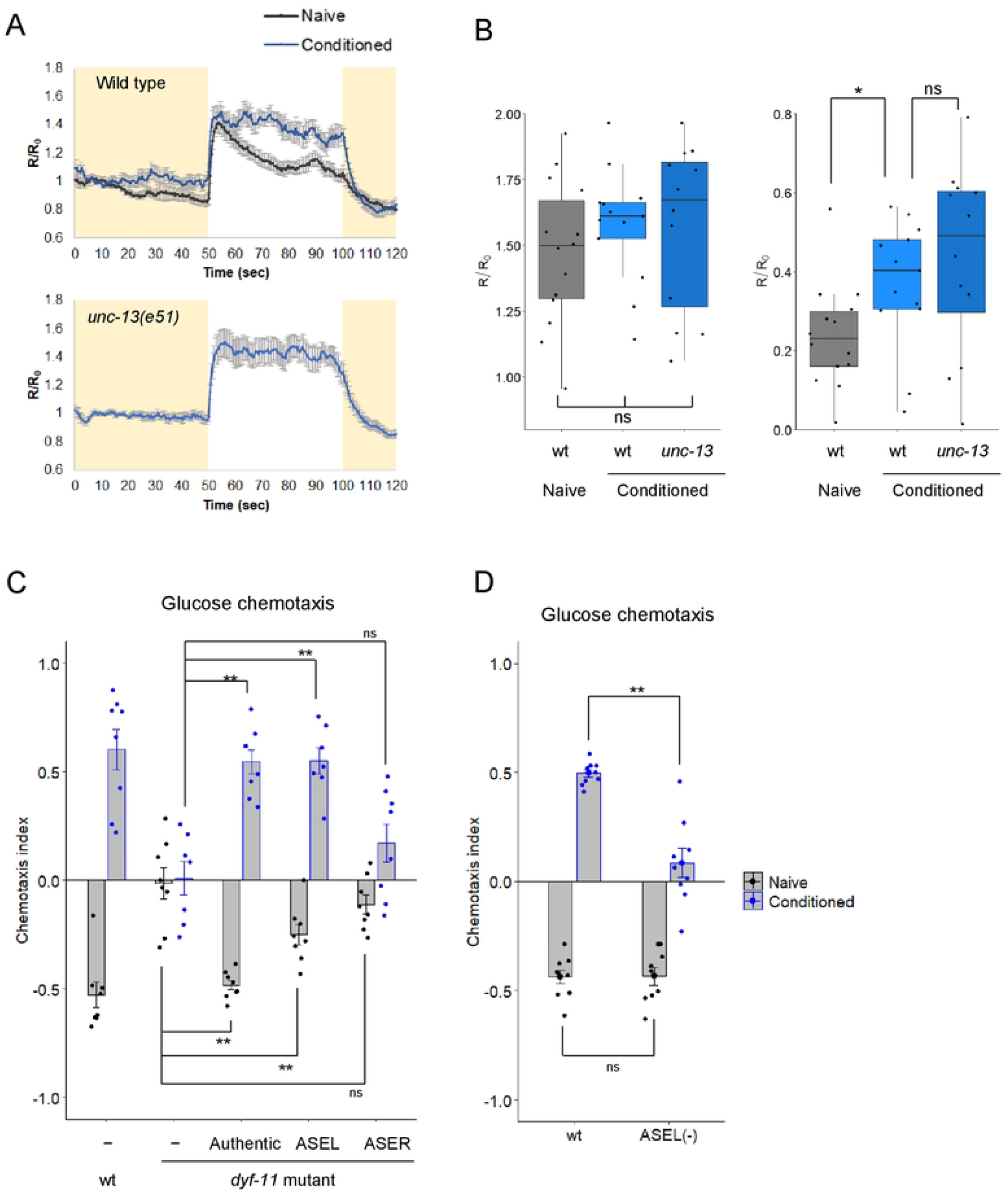
The ASEL neuron functions in glucose attraction after glucose conditioning. Calcium responses of ASEL when glucose concentration changes after conditioning with (“conditioned”) or without (“naïve”) glucose in the presence of food (A, B). (A) Time course of the average fluorescence intensity ratio (YFP/CFP) of YC2.60 relative to the basal ratio (R/R_0_) in ASEL. The glucose concentration was switched from 15 to 0 mM at 50 s and then returned to 15 mM at 100 s. (B) Quantification of calcium responses of ASEL upon decrease in glucose concentration. Peak fluorescence intensity ratios (R/R_0_) of YC2.60 in ASEL during the 10-s period after the decrease in glucose concentration were plotted (left). Differences between the average fluorescence intensity ratios (R/R0) within 0–50 s and 50– 100 s were plotted. Calcium responses in the wild type (naïve, n = 14; conditioned, n = 13) and *unc-13(e51)* (conditioned, n = 12). Two-tailed Welch’s *t*-test: \**P* < 0.05. Box plots represent the median (central line), 25th and 75th percentiles (the box), and the smallest and largest values that are not outliers (the whiskers). Individual data points are superimposed on the box plots. (C, D) After feeding conditioning with (conditioned) or without (naïve) glucose, chemotaxis to glucose was tested. Wild type without transgene and *dyf-11(pe554)* mutants expressing *dyf-11(+)* cDNA under the authentic promoter or the ASEL- or ASER-selective promoter or without the transgene were tested (C). Wild type (“wt”) and strain OH8585 (“ASEL (−)”), in which ASEL is genetically ablated, were tested (D). ANOVA followed by Dunnett’s *post hoc* test (C) or two-tailed Welch’s *t*-test (D): \*\**P* < 0.01. n = 8 (C) or 9 (D) assays.

We next examined glucose chemotaxis using the *dyf-11* mutant, in which sensation to water-soluble chemicals is largely disrupted because of sensory cilium defects. These defects can be recovered in a cell-type-specific manner by expression of *dyf-11*(+) cDNA in restricted cell types [8]. The *dyf-11* mutants showed no chemotactic response to glucose even after glucose conditioning. The glucose chemotaxis defects were significantly recovered by *dyf-11* expression in ASEL but not in ASER (Fig. 2C). Furthermore, genetic ablation of ASEL using caspase [22] caused a significant defect in glucose attraction after glucose conditioning (Fig. 2D). These data suggest that the ASEL neuron plays a major role in glucose attraction after glucose conditioning.

### Behavioral output of ASEL activation is changed after glucose conditioning

Calcium imaging experiments suggested that ASEL is activated when glucose concentration is decreased with or without glucose conditioning (Fig. 2A, B). We next monitored worm movements following activation of ASEL using a multiworm-tracking system combined with an optogenetic tool [12]. When ASEL was activated by *Channelrhodopsin2*, the probability of forward locomotion significantly increased in worms without conditioning (Fig. 3A and B, gray). By contrast, it decreased in worms after glucose conditioning (Fig. 3A and B, blue). These data suggest that the behavioral output of ASEL activation is altered by glucose conditioning. We propose a model in which ASEL is activated upon decreased glucose concentration and promotes or suppresses forward movement in worms without conditioning or after glucose conditioning, respectively, thereby switching the response from negative to positive chemotaxis through glucose conditioning.

**Figure 3.**
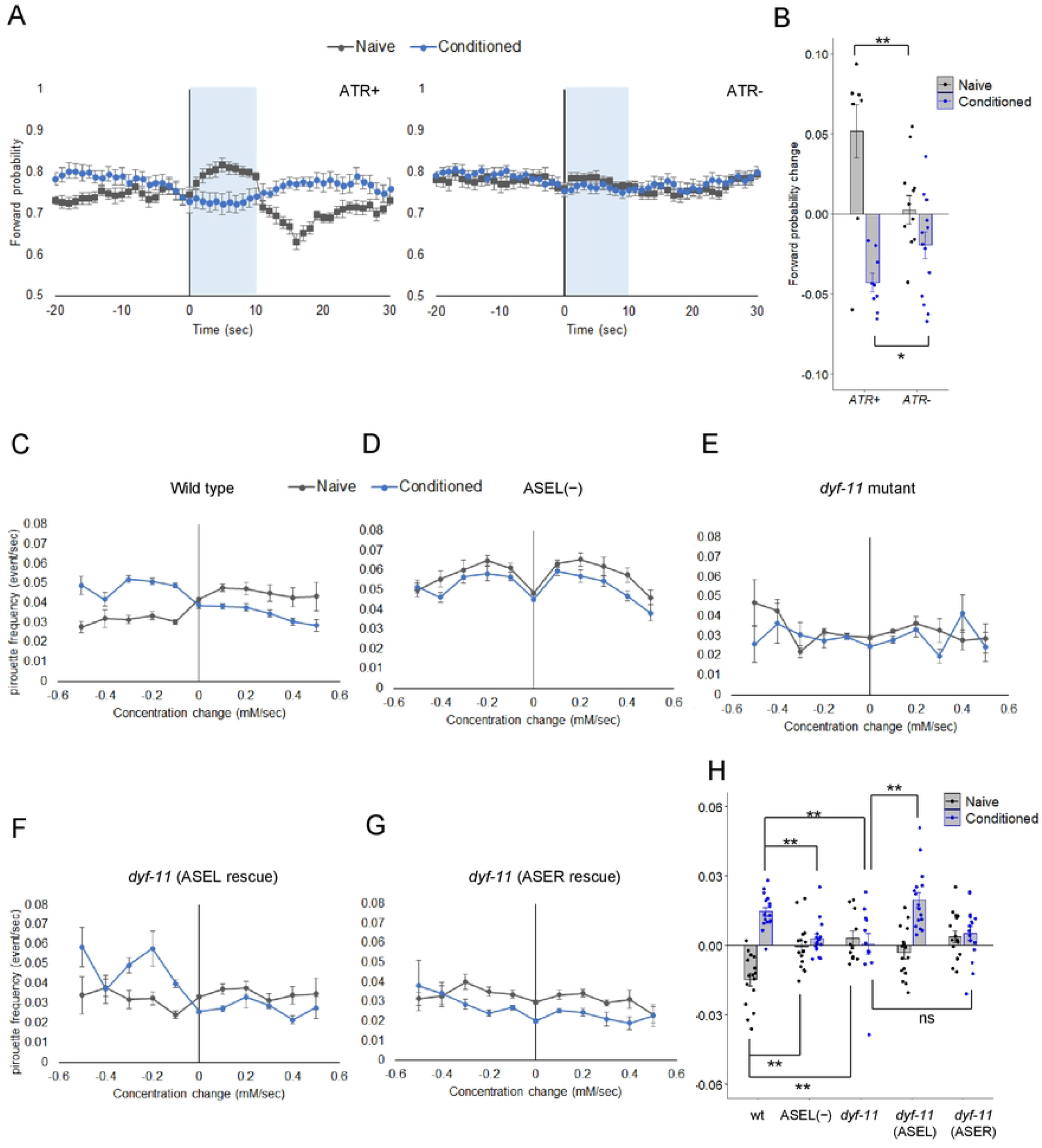
Behavioral outputs after glucose concentration changes are switched by glucose conditioning. Probability changes in forward movement by optogenetic activation of ASEL (A, B). (A) After feeding conditioning with (blue traces) or without (gray traces) glucose, probabilities of forward movement were monitored in worms expressing *ChR2* in ASEL on agar plates, containing 5 mM glucose. The ASEL neuron was activated by blue light illumination for 10 s (shaded in blue). To activate *ChR2*, all-*trans* retinal (ATR) was applied during conditioning (left). The same procedure was used without ATR as a control (right). (B) Changes in forward probability were quantified by subtracting average forward probability before photostimulation (−20–0 s) from that during photostimulation (3–7 s). Bars represent mean values; error bars represent SEM. n = 11–15. Two-tailed Welch’s *t*-test: \**P* < 0.05, \*\**P* < 0.01. Changes in pirouette frequency according to glucose concentration changes (C-G). After feeding conditioning with (blue traces) or without (gray traces) glucose, pirouette frequencies were monitored in the wild-type (C), the OH9019 ASEL-ablated strain (D), *dyf-11(pe554)* (E), *dyf-11(pe554)* strains expressing *dyf-11*(+) in ASEL (F), or ASER (G). Migration bias was quantified by using the pirouette index, defined as the difference of probability of pirouette between negative dC/dT rank and positive dC/dT rank. Positive and negative pirouette indices mean migration bias toward high and low glucose concentrations, respectively (H). Bars represent mean values; error bars represent SEM. n = 11–16. ANOVA followed by Tukey’s *post hoc* test: \*\**P* < 0.01.

A study reported that in *C. elegans*, chemotaxis is achieved by changing the turning rate as a function of concentration change [23]. We monitored tracks of worms during glucose chemotaxis using a multitracking system [24]. Each track was divided into periods of smooth swimming (runs) and frequent turning (pirouettes), based on recently defined criteria, and the frequency of pirouettes during glucose chemotaxis was determined [8, 23]. Pirouette frequency increased and decreased after increasing and decreasing glucose concentration, respectively, in wild type worms without conditioning (Fig. 3C and H first block, naïve). By contrast, those responses were reversed after glucose conditioning (Fig. 3C and H first block, conditioned). These changes in pirouette responses to glucose led to avoidance or attraction of glucose in worms without conditioning or those after glucose conditioning, respectively. Both ASEL-ablation and *dyf-11* mutant worms did not show significant changes in pirouette frequencies during glucose chemotaxis (Fig. 3D, E, H second and third blocks). The expression of *dyf-11* in ASEL, but not ASER, recovered increase in pirouette frequency in response to decreased glucose concentration after glucose conditioning (Fig. 3F, G, H fourth and fifth blocks, conditioned). These data are consistent with the notion that ASEL action is required and sufficient for glucose attraction after glucose conditioning.

### NaCl and glucose sensation by ASEL during feeding reduce chemotaxis to glucose and Na^+^, respectively

We previously reported that *C. elegans* is attracted toward Na^+^ after NaCl conditioning in the presence of food [12]. ASEL responds and drives attraction to Na^+^ in a way different from glucose: ASEL is activated by increased Na^+^ concentration and promotes forward locomotion after NaCl conditioning, thereby driving attraction to Na^+^. We examined both glucose and Na^+^ chemotaxis after exposure to a mixture of glucose and NaCl in the presence of food. Glucose conditioning in the absence of NaCl promoted glucose attraction, and interestingly, caused strong Na^+^ avoidance (Fig. 4A left, first block, Fig. 4B). Glucose attraction switched to glucose avoidance in the presence of NaCl along with glucose during conditioning (Fig. 4A left, blue bars). Conversely, the Na^+^ avoidance decreased by addition of NaCl along with glucose during conditioning (Fig. 4A left, red bars). After NaCl conditioning in the absence of glucose, the worms showed glucose avoidance and Na^+^ attraction; however, these responses gradually decreased by glucose addition along with NaCl during conditioning (Fig. 4A right). These data suggest that sensing of NaCl and glucose during feeding, reduced glucose and Na^+^ chemotaxis, respectively, and the ratio of these chemicals during conditioning determines the direction and extent of chemotaxis to Na^+^ and glucose. To rule out that high osmolality and not the chemical properties of glucose affect Na^+^ avoidance after glucose conditioning, worms were exposed to sucrose in the presence of food, at the same concentration as glucose. Sucrose conditioning did not lead to significant Na^+^ avoidance, suggesting that high osmolality during conditioning is not sufficient for the induction of Na^+^ avoidance (Fig. 4B). We next examined glucose-induced Na^+^ avoidance using the sensory cilia-defective *dyf-11* mutant and the *dyf-11* mutant, in which sensory cilium is rescued only in ASEL. The *dyf-11* mutant did not show prominent Na^+^ chemotaxis even after NaCl and glucose conditioning (Fig., 4C, the second block). By contrast, recovery of the sensory cilium of ASEL was sufficient for attraction and avoidance of Na^+^ after NaCl and glucose conditioning, respectively, suggesting that sensory perception by ASEL is required for two directions of Na^+^ chemotaxis after NaCl and glucose conditioning (Fig. 4C, the fourth block). Unlike the wild type control, the *dyf-11* mutant whose sensory cilium of ASEL was recovered showed strong Na^+^ attraction even without NaCl conditioning. A similar phenotype was observed in worms whose neuronal identity of ASER is converted to ASEL identity (2-ASEL strain) [22, 25]. They showed strong attraction to Na^+^ with or without NaCl conditioning, and glucose conditioning caused Na^+^ avoidance (Fig. 4D). These data imply that ASER functions to reduce Na^+^ attraction in conditions without high NaCl exposure in the presence of food.

**Figure 4.**
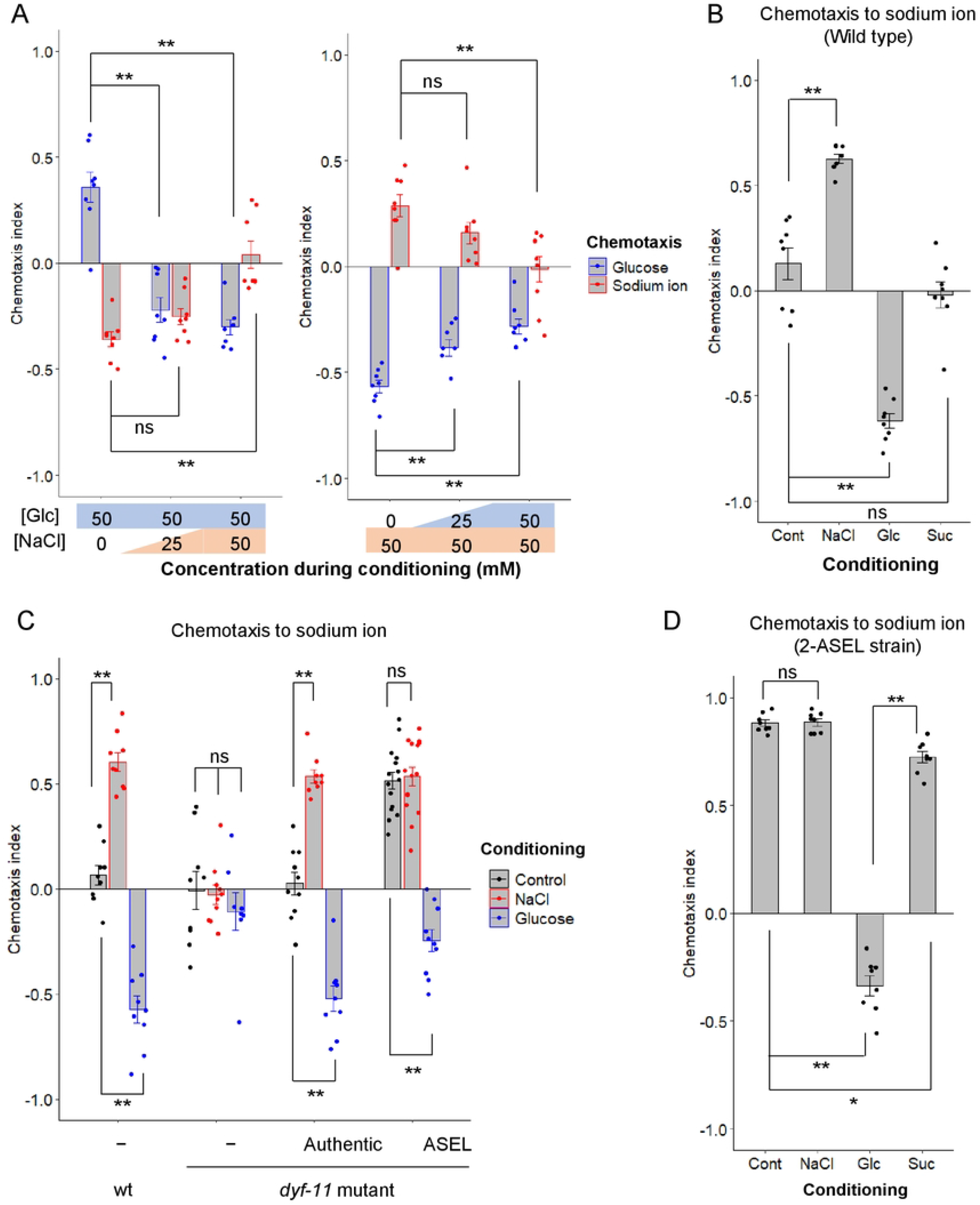
Conditioning with glucose and sodium ions decreases chemotaxis to sodium ions and glucose, respectively. After feeding conditioning with the indicated chemicals, chemotaxis to glucose (A, blue) or Na^+^ (A, red, B–D) was tested. 0, 25, or 50 mM NaCl and 50 mM glucose were used for conditioning (A, left); 0, 25, or 50 mM glucose and 50 mM NaCl were used for conditioning (A, right). The osmolarity was adjusted with sucrose to be the same in the conditioning plates in each experiment (A). Conditioning was performed with or without (“control” or “cont”) 100 mM of NaCl, glucose, or sucrose (B–D). The wild type was used (A and B). The wild type and *dyf-11(pe554)* mutants with or without transgenes expressing *dyf-11*(+) under the authentic promoter or the ASEL-selective promoter were used (C). The 2-ASEL strain, OH7621, was used (D). Bars represent mean values; error bars represent SEM. n = 8 (A), 6 (B, D), or 8–15 (C). ANOVA followed by Dunnett’s (A, C) or Tukey’s (B, D) *post hoc* test: \**P* < 0.05, \*\**P* < 0.01.

### PKC-1 promotes glucose attraction and Na^+^ avoidance after glucose conditioning

DAG/PKC-1 signaling plays significant roles in chemosensory neurons, including ASER gustatory and AWC olfactory neurons, to regulate chemotaxis plasticity [8, 26]. One of the functions of this signaling pathway in ASER is the promotion of chemotaxis toward high salt by increased neurotransmission from ASER to downstream interneurons [10, 11]. We examined glucose chemotaxis in a loss-of-function (lf) mutant of *pkc-1, pkc-1(nj3)*, which showed a strong defect in high-salt migration [8]. The *pkc-1(nj3)* mutant showed a strong defect in glucose attraction after glucose conditioning. Conversely, a gain-of-function (gf) mutant of *egl-30, egl-30(pe914)*, in which DAG/PKC-1 signaling is increased [27], showed attraction instead of avoidance without glucose conditioning (Fig. 5A). Weak glucose attraction was also observed by expression of a gf form of PKC-1 in ASEL, but not in ASER, without conditioning (Fig. 5B). These data suggest that PKC-1 in ASEL promotes glucose attraction after glucose conditioning. By contrast, the *pkc-1(nj3)* mutation promoted Na^+^ chemotaxis with or without glucose conditioning (Fig. 5C). The gf form of PKC-1 in ASEL or ASER reduced or increased Na^+^ chemotaxis, respectively (Fig. 5D). These data suggest that PKC-1 functions in opposite directions in ASEL and ASER and PKC-1 function is required for avoidance of Na^+^ after glucose conditioning.

**Figure 5.**
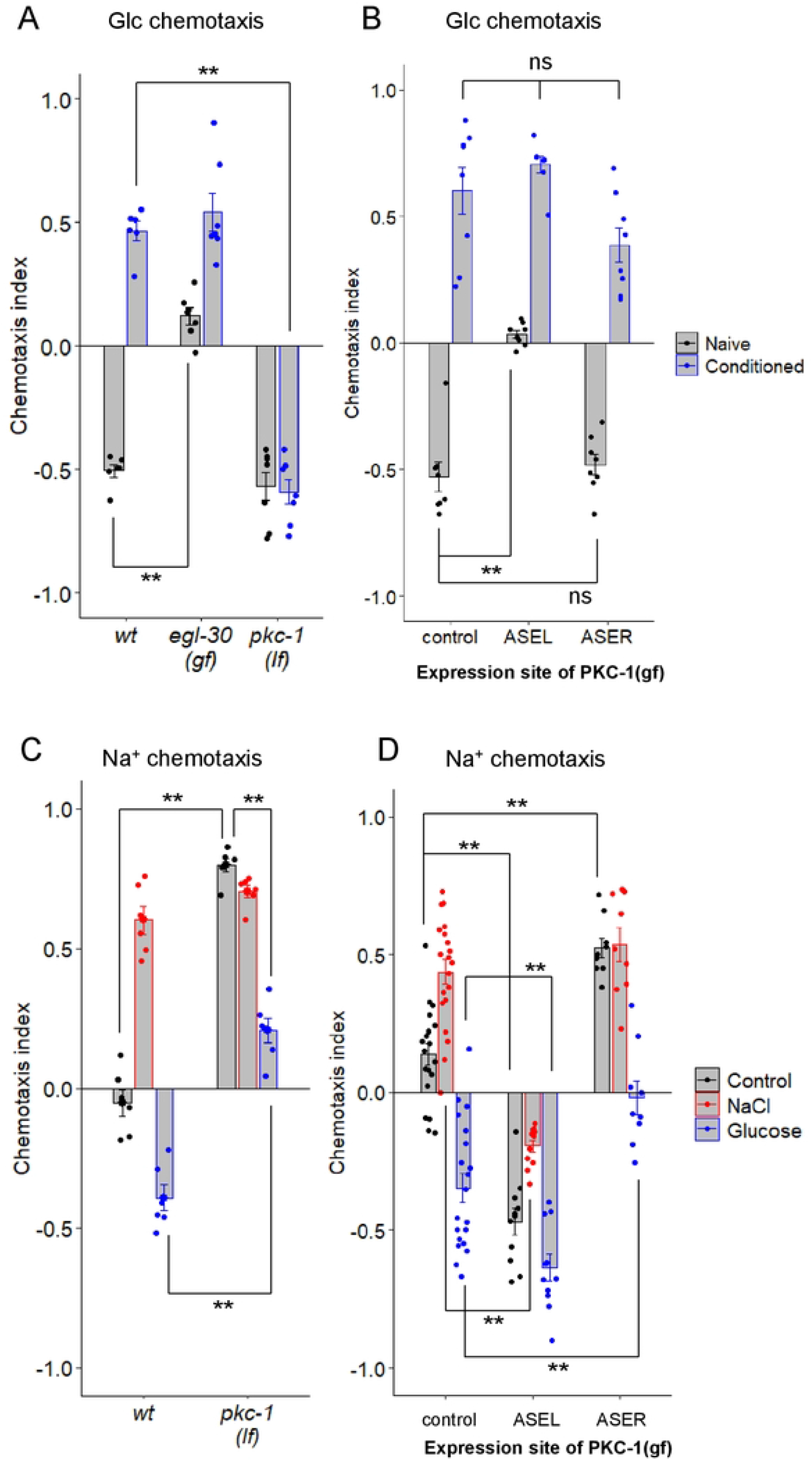
PKC-1 in ASEL promotes glucose attraction and Na^+^ avoidance after glucose conditioning. After feeding conditioning with (“conditioned”) or without (“naïve”) glucose, chemotaxis to glucose was tested (A, B). The wild type, *egl-30(pe914*gf) and *pkc-1(nj3*lf) (A) and wild type worms with or without (“control”) transgenes expressing PKC-1(A160E, gf) in ASEL or ASER (B). After feeding conditioning with or without (“control”) glucose or NaCl, chemotaxis to Na^+^ was tested (C, D). Wild type and *pkc-1(nj3*lf) (C) and wild type worms with or without (“control”) transgenes expressing PKC-1(A160E, gf) in ASEL or ASER (D). Bars represent mean values; error bars represent SEM. n = 6–7 (A), 8 (B), 6 (C), or 9–20 (D). ANOVA followed by Dunnett’s *post hoc* test (A, B, D) or two-tailed Welch’s *t*-test (C): \*\**P* < 0.01.

### ENSA-1 regulates chemotaxis after glucose conditioning in parallel with PKC-1

Although the *pkc-1(lf)* mutation promoted Na^+^ attraction in worms without conditioning, glucose conditioning significantly decreased Na^+^ chemotaxis (Fig. 5C). To further examine molecules required for Na^+^ chemotaxis plasticity after glucose conditioning, we performed forward genetic screening (see detail in the Methods section). We mutagenized the 2-ASEL strain, OH7621, and screened for mutants that showed Na^+^ attraction even after glucose conditioning. The isolated JN4784 showed strong Na^+^ attraction after glucose conditioning (Supplementary Fig. 3B). We identified a candidate region in which the causative mutation locates by a method based on whole-genome sequencing of the backcrossed progenies of JN4784 with either mutant or wild type phenotypes (see Methods). Within the candidate region, we found a nonsense mutation (Gln25*), *pe4796*, in *ensa-1*, which encodes an ortholog of the protein phosphatase inhibitor ARPP-16/19 (Supplementary Fig. 4B). Introduction of the fosmid WRM065cF04, which contains *ensa-1*, reduced increased Na^+^ attraction of the *ensa-1(pe4796)* mutant (Supplementary Fig. 3C). Two insertion/deletion mutants of *ensa-1, pe4791* and *pe4792*, which were obtained by using a CRISPR/Cas9-based method, showed increased Na^+^ attraction, similar to *ensa-1(pe4796)* (Fig. 6B). Furthermore, they showed defects in glucose attraction after glucose conditioning (Fig. 6A). Therefore, *ensa-1* is required for both glucose and Na^+^ chemotaxis plasticity after glucose conditioning, similar to *pkc-1*. Both *ensa-1* and *pkc-1* mutants showed significant increase in Na^+^ chemotaxis without conditioning compared with the wild type (Fig. 6B and C, black bars) and glucose conditioning significantly decreased Na^+^ chemotaxis of these mutants (Figs. 6B and C, compare black and blue bars). By contrast, in the *pkc-1; ensa-1* double mutant, glucose conditioning had no effect on Na^+^ chemotaxis (Fig. 6C second block, compare black and blue bars), suggesting the redundant function of *pkc-1* and *ensa-1*. Expression of *ensa-1* cDNA only in ASEL significantly rescued both glucose attraction and Na^+^ avoidance after glucose conditioning (Fig. 6D). Altogether, these data suggest that *ensa-1* functions in ASEL to promote glucose conditioning in parallel with *pkc-1*. PKC-1 is strongly localized to the presynaptic region in the ASER neuron [9]. Unlike PKC-1, an ENSA-1::Venus fusion protein was distributed throughout the neuron, including the cell body, axon, and dendrite, implying that ENSA-1 could function with PKC-1 in the presynaptic region and other subcellular sites in ASEL after glucose conditioning (Fig. 6E).

**Figure 6.**
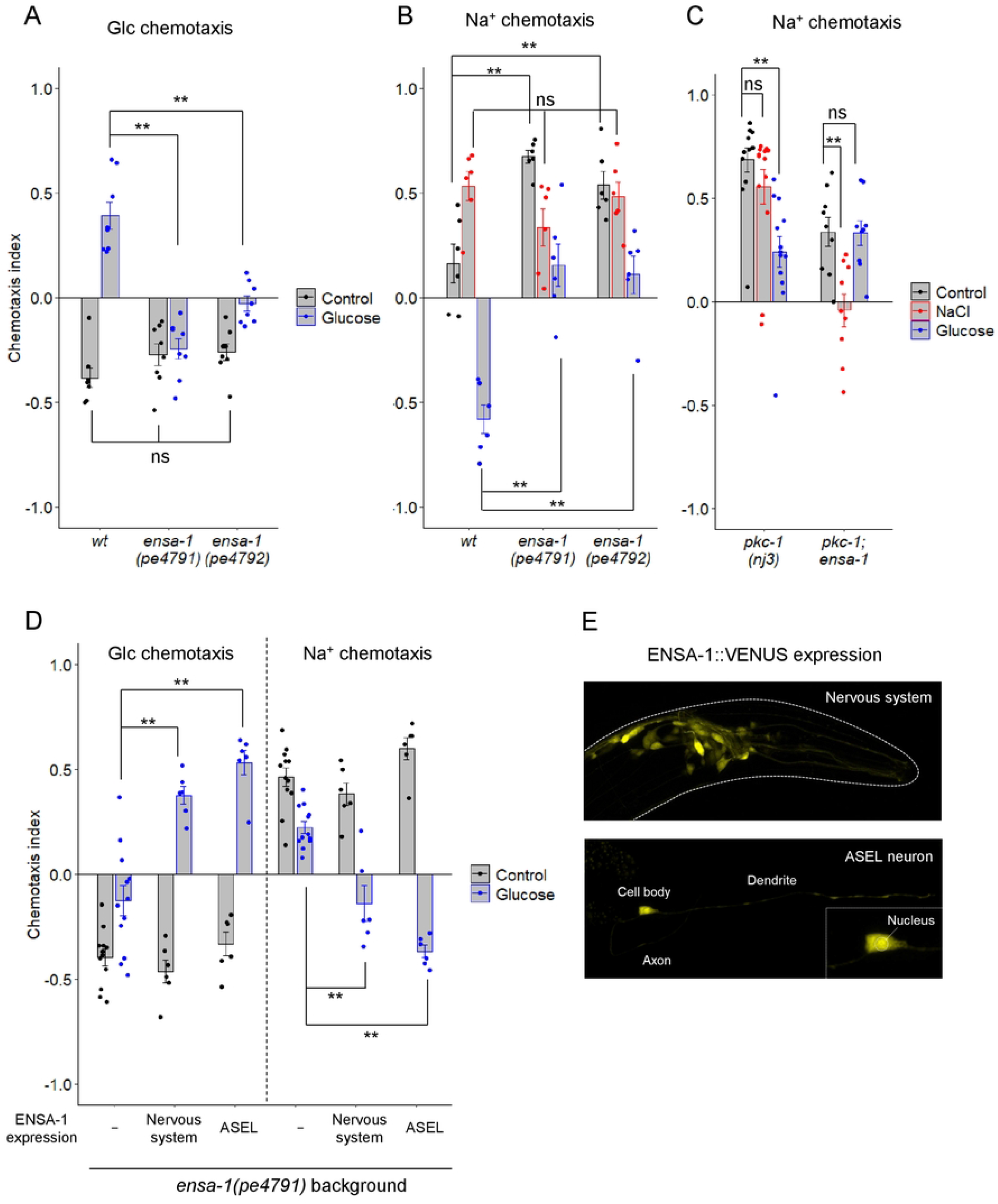
ENSA-1 in ASEL promotes glucose attraction and Na^+^ avoidance after glucose conditioning. (A-D) After feeding conditioning with or without (“control”) the indicated chemicals, chemotaxis to glucose or Na^+^ was tested. Bars represent mean values; error bars represent SEM. n = 8 (A), 6 (B), 7–9 (C), or 12 (D). ANOVA followed by Dunnett’s *post hoc* test: \*\**P* < 0.01. (E) Expression patterns of ENSA-1::Venus driven by the pan-neuronal *rgef-1* promoter (top) or the ASEL-selective *gcy-7* promoter (bottom). Head regions of adult worms are shown. An enlarged image of the cell body of ASEL is shown on the lower right.

## Discussion

*C. elegans* memorizes environmental signals, such as temperature, odorants, and salts, during feeding and learns to be attracted to those signals [8, 28, 29]. This type of behavioral plasticity may increase their chances of survival by moving toward an environment providing food. In this study, we report that worms avoid glucose and fructose; however, they learn to be attracted to the monosaccharides after cultivation with high concentrations of the monosaccharides in the presence of food. We found that the ASEL neuron is activated by decreased glucose concentrations and is required for glucose attraction. ASEL activation was reported to promote forward movements [12, 19]. We confirmed that optogenetic stimulation of ASEL promotes forward moving probability, thereby promoting glucose avoidance in response to decreased glucose concentration. By contrast, after glucose conditioning, ASEL activation decreased forward movements, suggesting that glucose conditioning alters the action of neural circuits downstream of ASEL and switches chemotaxis from avoidance to attraction of glucose. This phenomenon is reminiscent of plastic changes in neural circuits downstream of ASER in chemotaxis plasticity after high NaCl conditioning in the presence of food—high NaCl conditioning alters excitatory synaptic transmission from ASER to AIB interneurons, thereby promoting high NaCl attraction [10, 11]. ASEL may promote glucose attraction after high-glucose conditioning in the presence of food through a mechanism similar to that used in the ASER neural circuit. A similar mechanism acting in different neural circuits could contribute to robustness of food-seeking behavior based on food-associated learning to multiple chemicals, such as salts and monosaccharides.

The ASEL neuron has been shown to respond to several ions, including sodium, lithium, and magnesium [22]. ASEL responds to increase in concentrations of these ions. We have reported that ASEL promotes attraction to Na^+^ after conditioning with Na^+^ in the presence of food [12]. In this study, we observed the negative effects on chemotaxis to glucose and Na^+^ after Na^+^ and glucose conditioning, respectively. In addition to attractive responses to chemicals exposed during feeding conditioning, aversion of chemicals different from those exposed to during conditioning may promote food-seeking behavior by increasing accuracy to arrive at the chemical composition at conditioning. Glucose conditioning reduced forward moving probability during optogenetic ASEL activation. Because ASEL is activated by reduced glucose concentrations, ASEL activation following reduction in glucose would reduce forward movement, thereby promoting glucose attraction after glucose conditioning. Given that ASEL is activated by an increase in Na^+^ concentration [12], it is expected that ASEL activation after increasing Na^+^ concentration would reduce forward movement, thereby promoting aversion of Na^+^ after glucose conditioning. Therefore, activation of ASEL to concentration changes of glucose and Na^+^ in a opposite direction may underlie behavioral responses to those chemicals in the opposite direction after glucose conditioning. *C. elegans* AWC olfactory neurons were reported to change the response patterns to bacterially produced medium-chain alcohols in the context presenting more attractive odorants sensed by AWC, thereby switching from attraction to avoidance of medium-chain alcohols [30]. The chemosensory neurons of *C. elegans* can drive multiple sensory outputs according to the situation to survive.

DAG-PKC signaling has been identified as an important signaling pathway that regulates the direction and strength of taxis responses to temperature, salt, and odor in *C. elegans* [8, 26, 31]. PKC-1 promotes excitatory synaptic transmission from ASER to AIB interneurons to promote high NaCl attraction after high NaCl conditioning [10, 11]. Moreover, it promotes glutamatergic neurotransmission from AWC olfactory neurons to promote odorant attraction [32, 33]. Here, we report that chemotaxis plasticity after glucose conditioning depends on PKC-1 in ASEL. Although the activities of interneurons downstream of ASEL warrant further investigation, glucose conditioning may alter properties of neurotransmission from ASEL to downstream interneurons through DAG-PKC signaling in ASEL similar to that in ASER after high NaCl conditioning. Our forward genetic screening approach identified ENSA-1 (ARPP-16/19 ortholog), which regulates chemotaxis after glucose conditioning in parallel with PKC-1. ARPP-16/19 acts as an inhibitor of protein phosphatase 2A, PP2A, and is regulated by phosphorylation through MAST3 and PKA kinases [34]. Interestingly, KIN-4, the MAST3 kinase ortholog, regulates thermotaxis plasticity by regulating synaptic transmission from AFD thermosensory neurons to AIY interneurons [35]. ENSA-1, which distributes throughout the ASEL neuron, could interact with a variety of components of signaling pathways. It will be interesting to study how molecular signaling through PKC-1 and ENSA-1 regulates chemotaxis after glucose conditioning in ASEL.

## Materials and Methods

### Maintenance of *C. elegans* strains

*C. elegans* Bristol strain N2 was used as the wild type. The *C. elegans* strains were grown and maintained on NGM plates [36] seeded with *Escherichia coli* strain NA22 at 20°C, except for in calcium imaging and optogenetics experiments, in which *E. coli* strain OP50 was used as a food source. *C. elegans* strains used in this study are listed in S1 Table.

### DNA constructs and transgenesis

We used GATEWAY cloning (ThermoFisher) to generate plasmids for *YC2*.*60, ensa-1*, or *ensa-1::venus* expression. We generated the destination vector carrying the sequence of *YC2*.*60, ensa-1*, or *ensa-1::venus*, namely pDEST-YC2.60, pDEST-ensa-1::sl2::cfp, or pDEST-ensa-1::Venus::sl2::cfp, respectively, and then inserted the *rgef-1, gcy-5*, or *gcy-7* promoter sequence upstream of each gene through the LR reaction with the entry vector carrying each promoter. Venus was fused to the C-terminal region of *ensa-1* cDNA just before the stop codon by a PCR-based method. DNA constructs were injected at concentrations of 10–30 ng/µL with a co-injection marker, i.e., *lin-44p::mCherry* (20 ng/µL), *unc-122p::mCherry* (20 ng/µL), or *myo-3p::Venus* (10 or 20 ng/µL), and a carrier DNA, namely pPD49.26. Injection mixtures were prepared to a final concentration of 100 ng/µL.

For isolating *ensa-1(pe4791)* and *ensa-1(pe4792)* insertion/deletion mutants using a CRISPR/Cas9 system [37], we injected two plasmid DNAs, including *Cas9* cDNA under the *rgef-1* promoter or *ensa-1* sgRNA under the *U6* promoter, at 30 or 50 ng/µL, respectively, with a co-injection marker, namely *myo-3p::Venus* (10 ng/µL), into wild type N2. The target sequence (AGAGCTTATGGGCAAATTGG) in *ensa-1* sgRNA includes the first exon of *ensa-1*. We screened F1 animals harboring deletions or insertions around the *ensa-1* target sequence by PCR-based genotyping, using the MultiNA microtip electrophoresis system (Shimadzu). F2 animals harboring frameshift mutations in the first exon of *ensa-1* on two homologous chromosomes were isolated and mutation sites were determined.

### Behavioral assays

In all behavioral assays, we handled with chemotaxis buffer, which contains 25 mM potassium phosphate (pH 6.0), 1 mM CaCl_2_, and 1 mM MgSO_4_. For chemotaxis assays, except for worm-tracking experiments (Fig. 3A-F) and optogenetics experiments (Fig. 3G, H), we used circular agar plates (approx. 85 mm in diameter) with chemical gradients (Supplementary Fig. 1A, shown as chemotaxis-test plates). Chemical gradients on the test plates were produced as described [8, 12]. For chemotaxis to sugars (d-glucose, d-fructose, and sucrose), two cylindrical 2% agar blocks, including 0 mM or 50 mM sugar in chemotaxis buffer, were placed 3 cm from the center of a 2% agar plate, including the chemotaxis buffer, in opposite directions for 23–25 h at 20°C and removed just before the assay. For chemotaxis to Na^+^, two cylindrical 2% agar blocks, including 50 mM NH_4_Cl or NaCl in chemotaxis buffer, were placed 3 cm from the center of a 2% agar plate, including 50 mM NH_4_Cl in chemotaxis buffer, in the opposite directions for 18–24 h at 20°C and removed just before the assay. For conditioning, adult worms were transferred to modified NGM plates, including 100 mM d-glucose, d-fructose, sucrose, or NaCl instead of ∼51 mM NaCl in standard NGM, unless otherwise noted, for 4–5 h at 20°C. NaCl-free NGM plates were used for negative control (shown as “naïve” or “control”). *E. coli* NA22 cultured in NaCl-free LB liquid was seeded on the conditioning plates as a food source. After conditioning, worms were placed at the center of the chemotaxis-test plate and allowed to crawl for 45 min. Fifteen to two hundred worms were used in each assay. The chemotaxis index was determined according to the equation shown in Supplementary Fig. 1A.

For worm-tracking experiments, we used a rectangular agar plate (diameter approx. 78.5 mm × 121 mm) as a locomotion-test plate (Supplementary Fig. 1B). To form a linear glucose concentration gradient, a rectangular 2% agar block, including 50 mM glucose, was placed on a quartered area of a 2% agar plate for 23–25 h at 20°C and removed just before the assay. For glucose conditioning, adult worms were transferred to a modified NGM plate, including 100 mM d-glucose instead of ∼51 mM NaCl in standard NGM, for 4–6 h at 20°C. *E. coli* NA22 cultured in NaCl-free LB liquid was seeded on the conditioning plate as a food source. After conditioning, 30–50 worms were placed at the center of the test plate and recorded at one frame per second for 10 min using a multiworm-tracking system [24]. The analysis of tracking data was performed as reported [38]. The binarized images were used to extract the center-of-gravity coordinates of the worms. The trajectories of the extracted center-of-gravity coordinates with a length of 1 mm or longer were used as the worm’s trajectories in subsequent data analysis. To determine the concentrations of glucose on the test plate, 15 cylindrical agar blocks were excised from the central region of the test plate (Supplementary Fig. 1B) and the glucose concentrations were measured using Amplex Red Glucose/Glucose Oxidase Assay Kit (ThermoFisher). The cubic spline curve was created based on the average of three trials and used for subsequent locomotion analyses (Supplementary Fig. 1C). According to the definition in [38], periods of pirouette, consecutive sharp turns separated by less than 3.18 s, at all time points were determined and the frequencies of pirouette were calculated according to the time derivative of glucose concentration by 0.1 mM s^-1^ bins. Bins with <25 data points were omitted from further analysis. Pirouette index was defined as the difference of probability of pirouette between negative dC/dT rank and positive dC/dT rank.

Optogenetics experiments were performed as described [12]. We used a circular agar plate (approx. 85 mm in diameter), including 2% agar and 5 mM d-glucose in chemotaxis buffer, as a locomotion-test plate. We used transgenic worms expressing *Channelrhodopsin2* in ASEL in the mutant background *lite-1*, which encodes a blue light photoreceptor. For glucose conditioning, adult worms were transferred to a modified NGM plate, including 10 µM of all-*trans* retinal (ATR) and 0 mM or 100 mM d-glucose instead of ∼51 mM NaCl in standard NGM, overnight at 20°C. *E. coli* OP50 cultured in NaCl-free LB liquid was seeded on the conditioning plate as a food source. The modified NGM plate without ATR was used as negative control. After conditioning was complete, ∼50 worms were placed at the center of the test plate and recorded at one frame per second for 10 min using a multiworm-tracking system [24]. Blue light (peak wavelength = 470 mm; 0.2 mW/mm^2^) was delivered by a ring-shaped light emitting diode after 100 s of recording without light. For each experiment, light pulses of 10 s illumination were applied five times, with 80 s intervals between each pulse. Forward movement probability was calculated as the ratio of worms during forward locomotion, i.e., except during pirouettes, sharp turns, or pauses, at each time point, and values of five trials were averaged in each experiment. At least 11 experiments were performed for each condition.

### Calcium imaging

Experiments were conducted as described [8]. We used YC2.60 as a calcium indicator. For conditioning, adult worms were transferred to a modified NGM plate, including 0 mM or 100 mM d-glucose instead of ∼51 mM NaCl in standard NGM, overnight at 20°C. *E. coli* OP50 cultured in NaCl-free LB liquid was seeded on the conditioning plate as a food source. After conditioning was complete, worms were physically immobilized in a microfluidic device and an imaging buffer (25 mM potassium phosphate [pH 6.0], 1 mM CaCl_2_, 1 mM MgSO_4_, and 0.02% gelatin; osmolarity was adjusted to 350 mOsm with glycerol) containing 15 mM d-glucose was delivered to the tip of the nose. The glucose concentration contained in the imaging buffer was changed from 15 to 0 mM and then recovered to 15 mM after 50 s. Fluorescence intensities of CFP and YFP were simultaneously monitored at a rate of two frames per second. Fluorescence intensities in the cell body of ASEL or ASER were analyzed with custom-made scripts using the ImageJ software. The average fluorescence intensity of the ratio of YFP to CFP between 5 and 15 s from the start of recording was set as R_0_, and the fluorescence intensity ratio of YFP to CFP relative to R_0_ (R/R_0_) was calculated for a time series of images.

### Forward genetic screening

To isolate mutants defective in chemotaxis plasticity after glucose conditioning, we used the 2-ASEL strain (OH7621) [22, 39]. OH7621 showed strong attraction to Na^+^ without conditioning; however, the attraction to Na^+^ was strongly reduced after glucose conditioning (Fig. 4D). OH7621 was mutagenized and the F1 offspring were divided into 21 pools of approximately 6,000 each. Approximately 6000 worms of the next generation in each pool were conditioned with 100 mM glucose for 4–6 h and subjected to a Na^+^ chemotaxis assay. Worms attracted to higher Na^+^ were collected and the next generation of worms was cultured until they reached adulthood. This process was repeated eight times (Supplementary Fig. 3A). Finally, we isolated 3 worms from each pool and tested Na^+^ chemotaxis of offspring of each worm after glucose conditioning. One of the isolated strains, JN4784, showed strong attraction to Na^+^ after glucose conditioning (Supplementary Fig. 3B).

### Genetic mapping

To identify the region containing the gene involved in Na^+^ chemotaxis defect after glucose conditioning in JN4784, we performed variant discovery mapping based on whole-genome sequencing of the backcrossed offspring of JN4784 [40]. The JN4784 mutant was backcrossed with the original strain, OH7621, and then 99 lines of F1 generation progeny were isolated. Those cross progenies were cultivated across generations and their Na^+^ chemotaxis was tested after glucose conditioning two times in the F6 generation (Supplementary Fig. 4A). The top 20 lines of the average chemotaxis index were defined as the mutant-phenotype population and the bottom 10 lines were defined as the wt-phenotype population. The genome of each population was extracted and the genome fragment library was prepared using KAPA HyperPlus Library Preparation Kit for Illumina (NIPPON Genetics). Whole-genome sequencing was performed by multiplex paired-end Illumina Hiseq X Ten sequencing (Macrogen). Variants were extracted by mapping to the *C. elegans* reference genome at 104.07 average depth of coverage. After subtracting variants already present in OH7621, the variants in each mutant and wild type phenotype group were plotted according to their genomic positions. We found a candidate region with a high density of variants with a percentage close to 100% in chromosome I only in the mutant phenotype population (Supplementary Fig. 4B).

### Confocal microscopy

Anesthetized worms at the adult stage were imaged on a 5% agarose pad. Z-series images (slice spacing of 1 μm) of the head regions were acquired with a Leica SP5 confocal microscope using a 63×/1.30 objective. Z-stack images were created with ImageJ software.

### Statistical analysis

Statistical analyses were performed using R3.6.1 (http://www.R-project.org/). For multiple comparisons, ANOVA followed by Dunnett’s or Tukey’s *post hoc* test or two-tailed Welch’s *t*-test with the Holm correction was performed as indicated in each figure caption. All analyses were performed with at least four biological replicates.

## Acknowledgments

We are grateful to members of Iino’s laboratory, especially Dr. Hirofumi Kunitomo and Dr. Yu Toyoshima, for sharing unpublished experimental protocols and results and computer codes for data analysis. We thank the *Caenorhabditis* Genetics Center and the National Bioresource Project (Japan) for strains.

## Supporting information captions

**S1 Table. List of *C. elegans* strains**.

**S1 Figure. Chemotaxis- and locomotion-test plates**.

(A) Schematic of a chemotaxis-test plate used for chemotaxis assays, except for tracking analyses of worm locomotion. (B) Schematic for a locomotion-test plate used for tracking analyses using a multiworm-tracking system. Circles represent areas excised for measurement of glucose concentrations. (C) Glucose concentrations relative to positions on the *y*-axis in a test plate shown in B. Each curve from the three trials and the average curve smoothed by a cubic spline method are shown.

**S2 Figure. Calcium imaging of ASER upon change in glucose concentration**. Calcium responses of ASER upon glucose concentration changes after conditioning with glucose in the presence of food. Time course of the average fluorescence intensity ratio (YFP/CFP) of YC2.60 relative to the basal ratio (R/R_0_) in AESR. The glucose concentration was switched from 15 to 0 mM at 50 s and then returned to 15 mM at 100 s.

**S3 Figure. Screening for mutants defective in Na^+^ avoidance after glucose conditioning**. (A) Schematic of screening for mutants defective in Na^+^ avoidance after glucose conditioning. See detail in the Methods section. (B, C) After feeding conditioning with (“glucose”) or without (“control”) glucose was complete, chemotaxis to Na^+^ was tested. Original (OH7621) and isolated mutant (JN4784) strains were used (B). The JN4796 strain, which was isolated by outcrossing JN4784 with the wild type N2, with (+) or without (−) the fosmid, WRM065cF04, including *ensa-1* gene. See exact genotypes in S1 Table. Bars represent mean values; error bars represent SEM. n = 8–12 (B), 12 (C). Two-tailed Welch’s t-test: \*\**P* < 0.01.

**S4 Figure. Genetic mapping of the causative genes of the JN4784 mutant**.

(A) After feeding conditioning with glucose, chemotaxis to Na^+^ was tested in JN4784, OH7621 and 99 of F6 cross progenies, which were isolated after crossing JN4784 with OH7621. Bars represent mean with SEM. n = 4 (JN4784, OH7621), 2 (cross progenies). (B) Variant frequencies in cross-progeny populations showing mutant (purple) or wild type (sky blue) phenotype. The horizontal axis represents physical positions on each chromosome.

